# Distribution of target-site resistance mechanisms to nicosulfuron and glyphosate in *Amaranthus palmeri* accessions from two different regions of Türkiye

**DOI:** 10.64898/2026.09.21.753262

**Authors:** Filiz Erbaş, Sara Álvarez-Rodríguez, Michael Ozolins, Eric L. Patterson

## Abstract

Palmer amaranth (*Amaranthus palmeri* S. Watson) is a highly competitive weed that has evolved resistance to several modes of action and has recently become a significant problem in Türkiye. This study was designed to determine the distribution of common target-site resistance (TSR) mechanisms to the acetolactate synthase (*ALS*)-inhibiting herbicide, nicosulfuron, and the 5-enolpyruvylshikimate-3-phosphate synthase (*EPSPS*)-inhibiting herbicide, glyphosate, in *A. palmeri* accessions collected from two agricultural regions of Türkiye; the Çukurova Region and the Gediz Basin. A total of 96 accessions were analyzed for *ALS* gene mutations and relative *EPSPS* gene copy number variation. Whole-plant responses were also evaluated based on relative dry weight following treatment with nicosulfuron and glyphosate applied at twice the field rate. The *ALS* mutations Pro197Ser, Trp574Leu, and Ser653Asn were far more prevalent in plants from the Çukurova Region than those from the Gediz Basin. Elevated relative *EPSPS* gene copy numbers were more common and higher in the Gediz Basin. Single rate herbicide screening experiments confirmed that accessions from the Çukurova Region exhibited higher dry weight response to nicosulfuron, whereas those from the Gediz Basin had higher to glyphosate Multivariable regression analyses revealed that nicosulfuron response was primarily associated with the Pro197Ser and Trp574Leu mutations, while glyphosate response was correlated with *EPSPS* gene copy number. These findings reveal distinct regional patterns of TSR mechanisms in *A. palmeri* in Türkiye that are consistent with differences in herbicide use and/or introduction history, but the roles of local selection, demographic processes, and gene flow remain unclear.

## Introduction

*A. palmeri* (*Amaranthus palmeri* S. Watson) is a summer annual weed native to North America that has now been identified in dozens of countries on 5 continents (EPPO, 2026) and it was recorded in the flora of Türkiye for the first time in 2016 (Eren et al. 2016). Studies have revealed that it’s abundance in Türkiye’s Gediz Basin, Çukurova Region, and Southeastern Anatolia Region (Sırrı 2022; Çatıkkaş 2024, Erbaş et al. 2024) and that, depending on the weed density, it can cause yield losses of up to 82% in soybean (Dalkılıç 2026), 79% in cotton (Erbaş et al. 2025a), 43% in corn (Erbaş et al. 2025b), 60% in sunflower, and 70% in tomato (Ülgen 2021) in Türkiye.

Known for its high seed yield, rapid growth, adaptability to diverse environmental conditions, and high competitiveness with other crops, *A. palmeri* is also known to have evolved resistance to many herbicide mode of action within the Herbicide Resistance Action Committee (HRAC), including Groups 2, 3, 4, 5, 9, 10, 12, 14, and 27 (Ward et al. 2013; Roberts and Florentine 2022). These resistant populations can be found in both the native and invaded range around the world (Souza et al. 2025; Heap 2026). Of all these resistance cases, resistance to the 5-enolpyruvylshikimate-3-phosphate synthase (*EPSPS*)-inhibiting herbicide glyphosate and the acetolactate synthase (*ALS*)-inhibiting herbicide nicosulfuron are the most concerning for Turkish farmers, as these are the most used herbicides in Türkiye (Torun 2017).

Herbicide resistance mechanisms are largely divided into two major categories: target-site resistance (TSR) mechanisms, and non-target site resistance (NTSR) mechanisms (HRAC 2026). The most prevalent TSR mechanism in *A. palmeri* is the amplification of the target gene *EPSPS* (Gaines et al. 2010; Singh et al. 2018; Molin et al. 2020). For *ALS* inhibitors, the most prevalent TSR mechanism in *A. palmeri* are point mutations in the *ALS* gene. These mutations reduce the binding affinity of *ALS*-inhibiting herbicides to the *ALS* protein, while preserving the functionality of the enzyme. The known *ALS* mutations in *A. palmeri* so far are Ala122Thr (A122T), Ala122Ser (A122S), Ala122Val (A122V), Pro197Ile (P197I), Pro197Ser (P197S), Pro197Ala (P197A), Pro197Arg (P197R), Pro197Thr (P197T), Ala205Val (A205V), Asp376Glu (D376E), Trp574Leu (W574L), and Ser653Asn (S653N) (Palmieri et al. 2022a; Manicardi et al. 2023; Kaya Altop et al. 2025; Ji et al. 2025; Trebol-Aizpurua et al. 2025; Heap 2026). NTSR resistance has also been described for *ALS* inhibitors via cytochrome P450 monooxygenases (P450s) detoxification in *A. palmeri* but seems to be far less common (Küpper et al. 2017; Nakka et al. 2017; Trebol-Aizpurua et al. 2025).

Recently, two-year pot studies were conducted in Türkiye to ascertain the efficacy of various herbicides at controlling *A. palmeri,* concluding that herbicides in HRAC Group 2 (foramsulfuron + iodosulfuron methyl-sodium, imazamox, rimsulfuron, and tribenuron methyl), Group 4 (2,4-D EHE), Group 5 (metribuzin and fluometuron), and mixed Group 27 + Group 2 (mesotrion + nicosulfuron) used in the early post-emergence period were ineffective in achieving adequate (> 90%) control (Çatıkkaş 2024). The ED_50_ values obtained in the dose-response studies conducted by Kaya Altop et al. (2025) also demonstrated that three Turkish populations of *A. palmeri* exhibited more than 7-fold resistance to glyphosate, 9.21–10.35-fold to nicosulfuron, and 6.41–7.44-fold to foramsulfuron + iodosulfuron methyl-sodium. These findings underscore the necessity to investigate the target-site resistance mechanisms for *ALS* and *EPSPS* inhibitors at the regional level in *A. palmeri*.

The objective of this study was to determine the presence of *ALS* mutations and relative *EPSPS* copy numbers in accessions collected from two regions in Türkiye where *A. palmeri* is present, and to assess dry weight response of plants at a 2x field rate. The study also aimed to characterize the distribution of these resistance-associated mechanisms among *A. palmeri* accessions from two regions.

## Materials and Methods

### Plant Material

*A. palmeri* seeds were collected in 2023 from 96 individual female plants located at least 5 km apart in two different agricultural regions of Türkiye: the Gediz Basin (western Türkiye) and the Çukurova Region (southern Türkiye). The information of seed collection sites is given in the Supplementary Table 1. Seed heads were dried for 7 days and manually cleaned by using sieves. Half of the seeds were sent from Türkiye to Michigan State University (USA) for molecular analysis. The remaining seeds were used in pot experiments at Aydın Adnan Menderes University (Türkiye) to ascertain the response of accessions to two-fold doses of nicosulfuron and glyphosate. Offspring of same female plants were used for molecular and pot studies.

### Genomic DNA Extraction

*A. palmeri* plants were grown in a greenhouse set at 30°C temperature, 40% humidity, 12/12 h day/night photoperiod supplemented with 278 µmol m ² s ¹ illumination by LED lighting. The seeds were sown in 20 cm diameter round pots filled with peat-based potting soil, and the seedlings were transplanted into 9 x 9 cm square pots containing the same substrate once they reached the 2-true leaf stage. Young leaf tissue of 8 plants for each offspring of same accession was collected in microcentrifuge tubes once plants reached the 6-8-leaf stage. These microcentrifuge tubes were then immediately transferred to tanks containing liquid nitrogen, and frozen leaves were ground inside the microcentrifuge tubes using an electric screwdriver with a pellet pestle attached to its end, and then stored in a freezer at-80 °C.

Genomic DNA was extracted from plant tissues using a modified CTAB protocol (Doyle and Doyle 1990). In brief, approximately 100 mg of frozen tissue was mixed with 850 µl CTAB buffer that contains 2% β-ME and incubated at 65 °C for 60 min. Following this, the microcentrifuge tubes were centrifuged at 15,000 rpm for 2 min. Thereafter, 1 µl of 10 mg ml^-1^ of RNase was added to each tube, and the tubes were placed into a dry block set at a temperature of 37 °C for 30 min. Following a 2 min centrifugation at 15,000 rpm, samples were extracted with 600 µl and 400 µl of chloroform:isoamyl alcohol (24:1), respectively. Each extraction was accompanied by a subsequent centrifugation process for 3 min, transferring the supernatant (600 µl and 450 µl, respectively) to a new tube. After adding 0.1 V 3 M NaOAc (∼ 450 µl) and 3.0 V pure cold EtOH (∼ 1350 µl), samples were centrifuged at 4 °C for 30 min. Following precipitation, DNA was washed with 400 µl of 70% cold ethanol. Excessive liquid poured slowly onto Kimwipes, prior to 10 min of evaporation. DNA was eluted in 35 µl of DNase/RNase free water and DNA concentration and quality was measured by using a NanoDrop spectrophotometer. Then samples were stored at-80 °C for further analysis.

### ALS Gene Sequencing and Genotyping Resistance Mutation

To amplify the *ALS* gene and cover all known mutation sites, several primer pairs were designed and tested. The primer pairs used in the analysis, together with the mutation sites covered and expected amplicon sizes, are presented in the Table 1. For each accession, amplification was initially attempted using F4/R4 primer pair, which covers all target mutation sites. If successful amplification was not obtained, additional primer combinations were tested sequentially. The F1/R3 and F2/R4 primer pairs were then used to amplify the containing P122 site and P197, A205, D376, W574 and S653 sites, respectively. If amplification remained unsuccessful, the F1/R3 and F2/R2 primer combination was tested, followed by the F5/R5 and F6/R6 primer pairs (Ji et al. 2025). This procedure was continued until successful amplification was obtained for at least three individuals per accession. Optimal annealing temperatures of all primers was 54 °C, which was determined by performing a temperature gradient PCR.

**Table 1.**
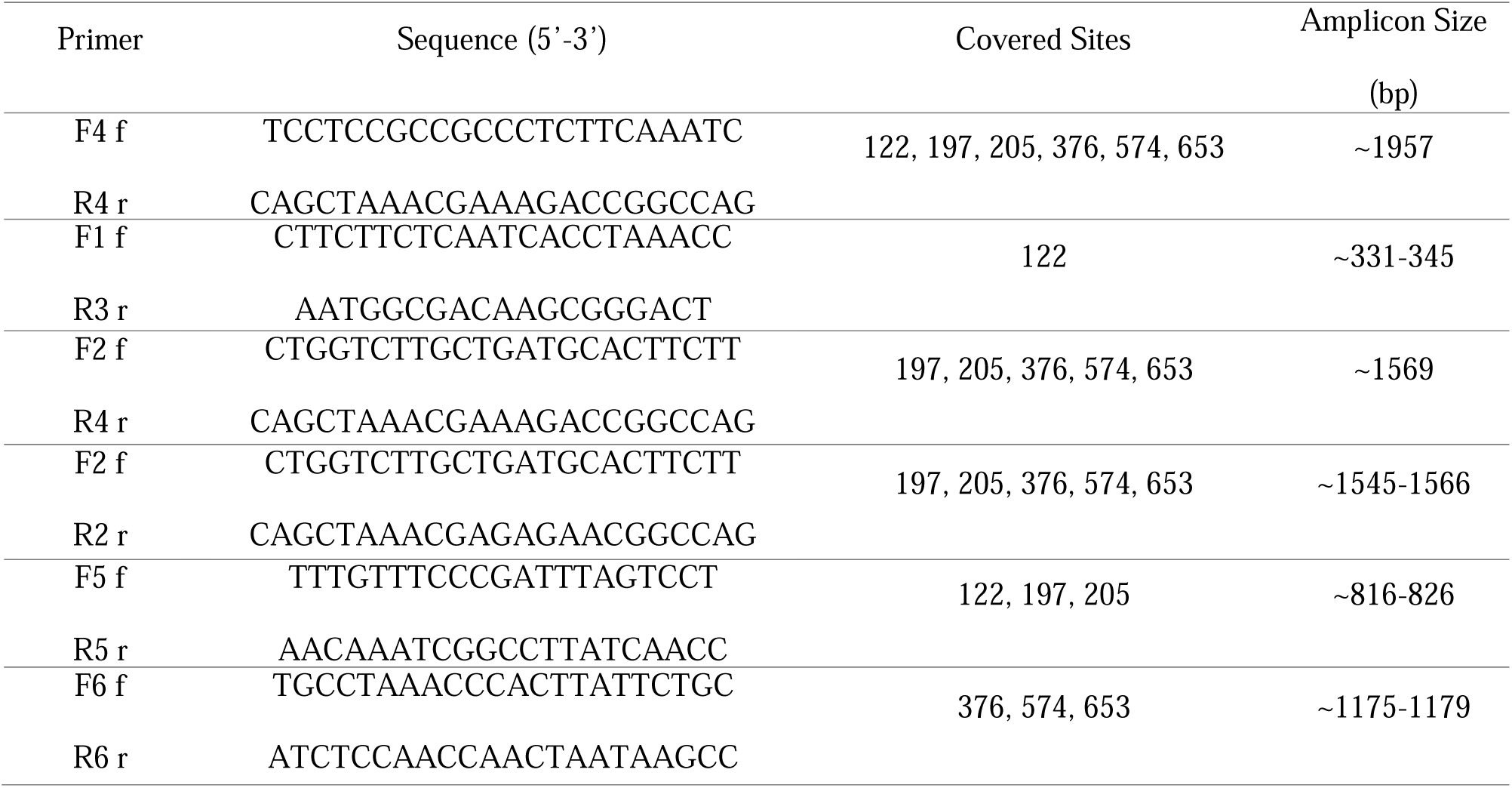
Primer pairs used for amplification of the *ALS* gene in Palmer amaranth (*Amaranthus palmeri*), including the mutation sites covered and the expected amplicon sizes.

The PCRs consisted of 12.5 μL GoTaq G2 Green Master Mix (Promega, 2800 Woods Hollow Road, Madison, WI 53711), 8.5 μL ultrapure water, 1 μL of forward and reverse primers (10 µM) and 2 μL gDNA to bring the total volume to 25 μL. The PCR was performed under the following conditions: initial denaturation at 95°C for 5 min, followed by 39 cycles of denaturation at 95°C for 30 s, annealing at 54°C for 30 s, an extension at 72°C for 2 min, and a final extension at 72°C for 8 min. The PCR tubes were stored at 4 °C until processed for sequencing. The presence of target fragments was confirmed by visualizing the PCR products on a 1% agarose gel. The PCR products were purified with Thermo Scientific GeneJet PCR Purification Kit and replicates were combined in equal proportions if obtained from the same offspring by using the same primers before nanopore sequencing. The results were analyzed in Excel files converted from fasta files. Frequency (*f*) of *ALS* mutations was calculated according to the following formula:

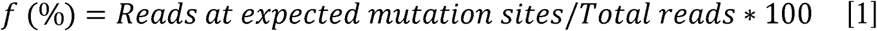

The average of *ALS* mutation frequencies across 3 replicates is presented in the results section.

### Relative EPSPS Gene Copy Number Determination

The relative gene copy number of the *EPSPS* gene was determined by quantitative real-time polymerase chain reaction (qPCR) in relation to the *A36* gene (Singh et al. 2018). The *A36* gene was used as a single copy reference. The amplification of the *A36* gene was achieved by employing the following set of primers: *A36*_F244 (5′-TTGGAACTGTCAGAGCAACC-3′) and *A36*_R363 (5′-GAACCCACTTCCACCAAAAC-3′), developed by Singh et al. (2018), while the amplification of the *EPSPS* gene was achieved by using the primer sets of EPSF1 (5′-ATGTTGGACGCTCTCAGAACTCTTGGT-3′) and EPSR8 (5′-TGAATTTCCTCCAGCAACGGCAA-3′), which were designed by Gaines et al. (2010).

For CNV qPCR, 10 µL of Bio-Rad IQ SYBR Green Supermix (Bio-Rad, 1000 Alfred Nobel Drive Hercules, California 94547), 6 µL of ultrapure water, 1 μL of the forward and reverse primers (10 µM) and 2 µL of gDNA were used to make the total volume of 20 µL. The real-time qPCR cycles started with 3 min at 95 °C, followed with 40 cycles of 95 °C for 30 s and 1 min at 60 °C. Melting curves were arranged from 65 °C to 95 °C with the increments of 0.5 °C for 5 s in a Bio-Rad CFX 96 real-time system thermocycler. A negative control consisting of primers with no template DNA was included in all reactions. The data were analyzed using a 2^ΔCt^ method to express the copy number of *EPSPS* relative to *A36*, as ΔCt= (Ct *A36* − Ct *EPSPS*). The relative increase in *EPSPS* copy number was expressed as 2^ΔCt^ (Gaines et al. 2011). Each accession sample was run in triplicate, and the average increase in relative *EPSPS* copy number is given in the results section.

### Evaluation of A. palmeri Response to Nicosulfuron and Glyphosate

These studies were conducted in the greenhouse of Aydın Adnan Menderes University of Türkiye between July 28, 2025 and November 4, 2025. Average air temperatures inside the greenhouse during the studies are shown in Supplementary Figure 1. During pot experiments, each offspring of same accession was subjected to an untreated control or a treatment with double dose glyphosate or nicosulfuron (2,880 g a.e. ha^-1^ for glyphosate (Ridown, 480 g ae L^-1^, SL, Rainbow) and 100 g a.i. ha^-1^ for nicosulfuron (Kalson, 40 g ai L^-1^, OD, Agrobest)). All herbicides were applied at 3-4 leaf stages of *A. palmeri* using a backpack sprayer equipped with flat fan nozzles at a pressure of 3 bars and a spray volume of 350 L ha^-1^. The plants were harvested above ground 21 days after treatment (DAT), left at 70°C for 48 h, and their dry weights were measured. Subsequently, the relative dry weight of each accession was calculated by comparing herbicide-treated plants with the mean dry weight of their respective control pots, according to the following formula:

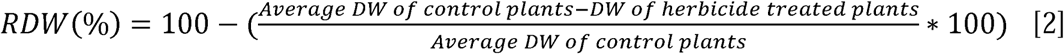

In the formula, RDW represents relative dry weight and DW represents dry weight. A completely randomized design was employed in two experimental runs, with 8 replicates in each one, by using a total of 4,608 pots filled with peat soil.

### Data Analysis and Mapping

General Linear Model (GLM) was performed using IBM SPSS Statistic v.21 to evaluate differences between dry weight responses of *A. palmeri* to double dose nicosulfuron and glyphosate applications by accepting region and experimental run as fixed factors. Since no significant main and interaction effects of experimental run were observed, the data were pooled and subsequently accession means analyzed to assess the relationship between relative dry weight and the presence of TSR mechanism. The analysis of potential variations in *ALS* mutation frequencies and relative *EPSPS* copy numbers across different regions was also conducted by incorporating the region as a fixed factor within a GLM. Univariable and multivariable linear regression analyses (enter model) were used to evaluate the associations between TSR mechanisms and relative dry weight of *A. palmeri* with/without addition of geographic region. As the different offspring from the same female plant were utilised in the determination of these associations, the identified associations remained at the family level. Information on *ALS* mutation frequencies and relative *EPSPS* copy numbers of *A. palmeri* accessions collected from Çukurova and Gediz Basins was mapped using QGIS 3.26.1 program.

## Results

### ALS Resistance Mutations

*ALS* sequences from all individuals were screened for mutations at six amino acid positions previously associated with resistance (A122, P197, A205, D376, W574, and S653). Among the specific substitutions examined, P197S, P197I, W574L, and S653N were detected in at least one individual, whereas A122S, A122T, A122V, P197A, P197R, P197T, A205V, and D376E were not detected in any individual more than 5%. The frequencies of *ALS* mutations detected based on *A. palmeri* accessions are given in (Figure 1-2 & Supplementary Table 2). In brief, P197S mutation was identified in 75% of the accessions in the Çukurova Region, while only in 6.3% of the accessions from the Gediz Basin (Figure 1A & Figure 2A). A single accession (GE-36) contained the P197I substitution in the Gediz Basin (Figure 2B). The mutations W574L and S653N were also prevalent in the Çukurova Region, appearing in 77% and 75% of the accessions, respectively (Figure 1B, C). These same mutations were also found in Gediz Basin, although at a lower frequency of 27% for both mutations (Figure 2C, D). Each accession collected from the Çukurova Region contained at least one *ALS* mutation with a frequency higher than 10%. However, in the Gediz Basin, 43.4% of the accessions had no *ALS* mutations (Figure 1-2 & Supplementary Table 2). Several *A. palmeri* accessions contained more than one *ALS* mutations (Figure 3). In the Çukurova Region, the P197S+W574L+S653N collection is the most common in 39.6% of accessions samples, while the P197S+W574L combination occurs in 20.8% of accessions. In the Gediz Basin region, the P197I+W574+S653N mutation, which is not found in the Çukurova Region, was only found in a single accession.

**Figure 1.**
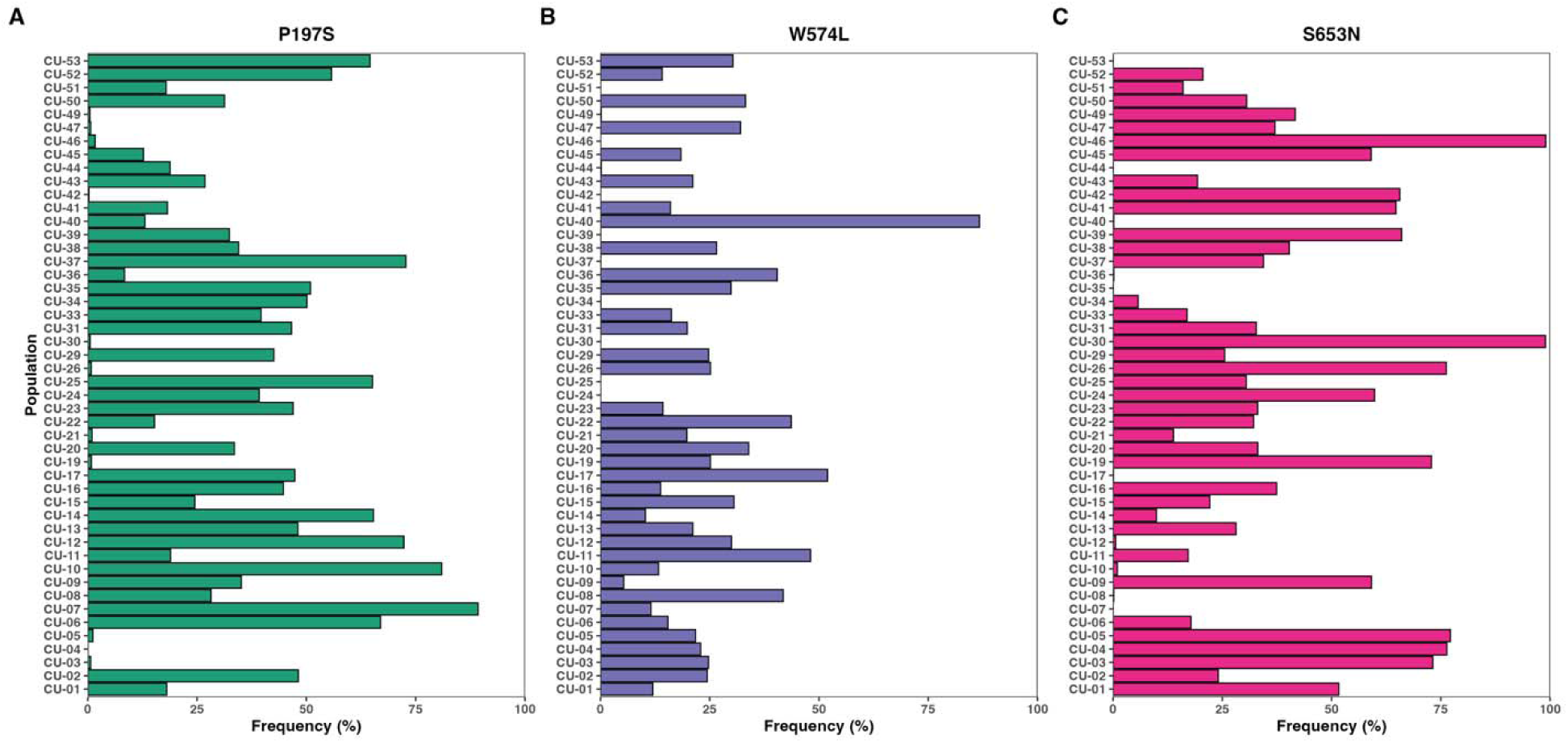
Mutation frequency of *ALS* mutations in Palmer amaranth accessions from the Çukurova Region (southern Türkiye). Mutation frequency (%) was determined by averaging three replicates. Panels represent individual specific mutations: (A) P197S (teal), (B) W574L (purple), and (C) S653N (pink). Bars indicate the mutation frequency for each studied accession.

**Figure 2.**
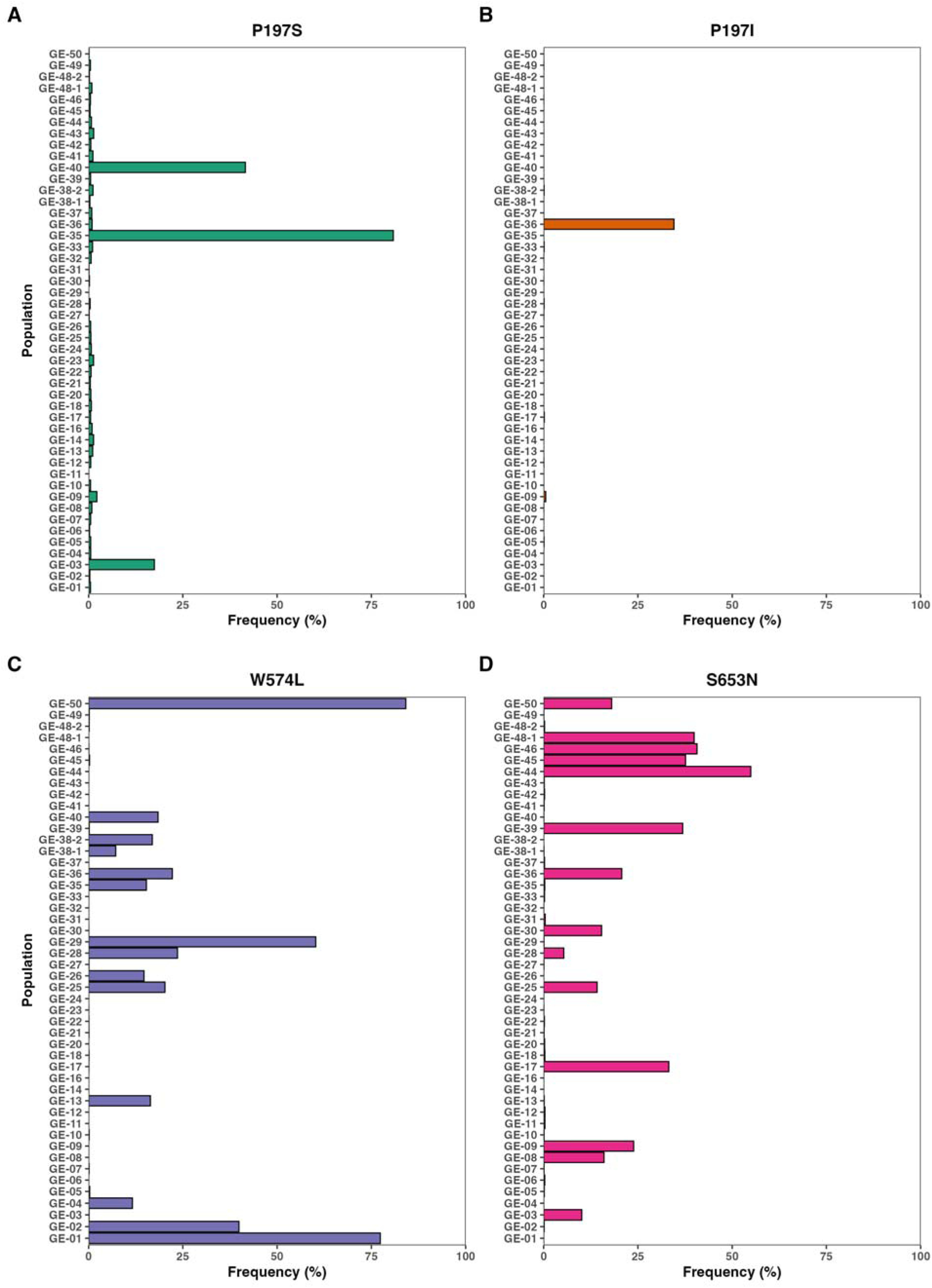
Mutation frequency of *ALS* mutations in Palmer amaranth accessions from the Gediz Basin (Türkiye). Mutation frequency (%) was determined by averaging three replicates. Panels represent individual specific mutations: (A) P197S (teal), (B) P197I (orange), (C) W574L (purple), and (D) S653N (pink). Bars indicate the mutation frequency for each studied accession.

**Figure 3.**
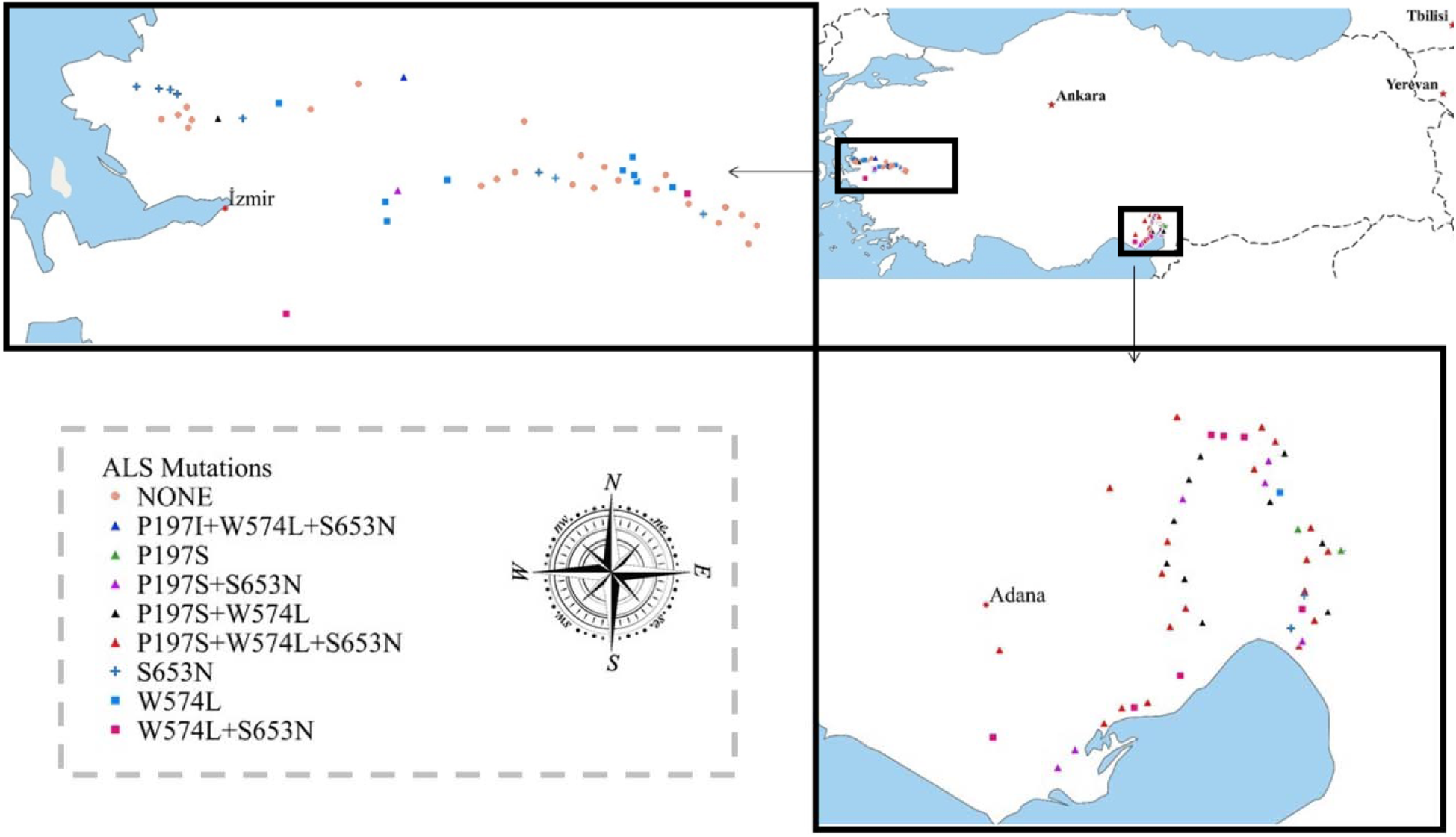
Geographic distribution and profile of *ALS* mutations in Palmer amaranth accession across Türkiye. The main map displays the surveyed collection sites, with zoomed-in insets highlighting the Gediz Basin (western Türkiye, top left) near İzmir and the Çukurova Region (southern Türkiye, bottom right) near Adana. Colored symbols represent the specific *ALS* target-site mutations or combination of mutations identified at each sampling location, as defined in the legend.

The statistical analyses given in Table 2 also confirmed that there are differences between the Çukurova Region and the Gediz Basin in terms of some *ALS* mutations. The mutation frequencies of P197S, W574L, and S653N appear to be higher in the Çukurova Region than in the Gediz Basin.

**Table 2.** Association between geographical region (Çukurova Region and Gediz Basin) and the frequency of *ALS* mutations. N: Number, p: Probability value.

| Factors | N | P197S | W574L | S653N |
| --- | --- | --- | --- | --- |
| Region |  | p=0.000 | p=0.003 | p=0.000 |
| Çukurova Region (CU) | 144 | 31.925±2.460 | 20.058±1.676 | 33.289±2.519 |
| Gediz Basin (GE) | 144 | 3.498±1.191 | 8.962±1.799 | 7.365±1.391 |

### Relative EPSPS Copy Number Variation

In the Çukurova Region, 27.1% of the accessions had just one relative copy of *EPSPS*, 60.4% of the accessions had between 2-40, 4.2% between 40-100, and 8.3% between 100-200 (Figure 4A, C & Supplementary Table 3). No accessions were found to have greater than 200 relative copies of *EPSPS.* In the Gediz Basin region, relative copy number variation was not detected in 12.5% of the accessions (Figure 4B, C & Supplementary Table 3). However, 41.7% of the accessions had between 2-40 relative copies of *EPSPS*, 25% had between 40-100 copies, and 12.5% between the 100-200 copies. Furthermore, in the Gediz Basin, 4 accessions contained relative *EPSPS* copy numbers estimated to be >200 (Figure 5). Relative *EPSPS* copy number has also been statistically determined to have significant between regions (p<0.001, N=288).

**Figure 4.**
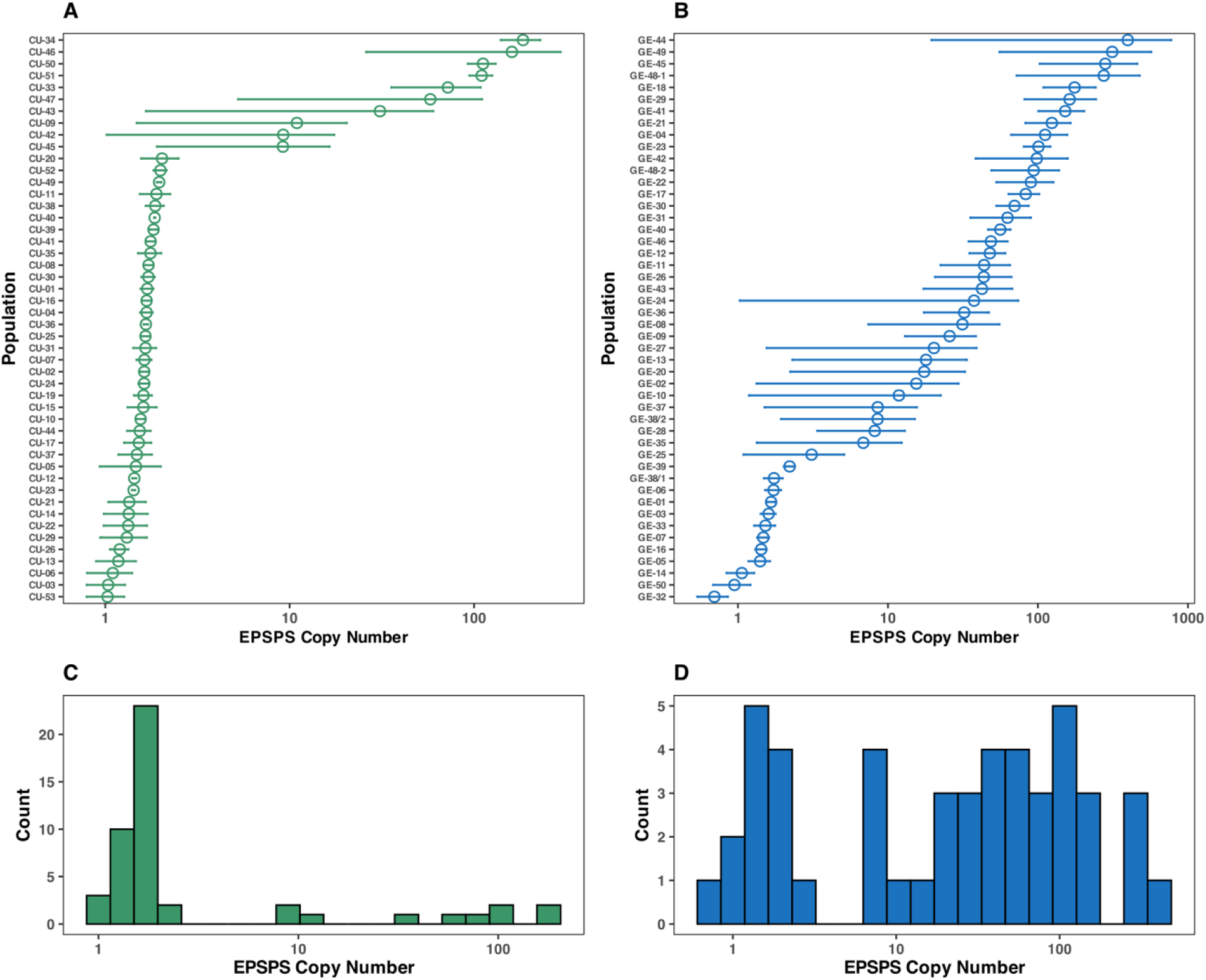
Variation in relative *EPSPS* copy number across Palmer amaranth accessions across Türkiye. (A–B) Mean relative *EPSPS* copy number (± SE) for Çukurova Region CU (A) and Gediz Basin GE (B). Horizontal error bars represent standard errors, and points indicat accession means. (C–D) Frequency distributions of relative *EPSPS* copy number corresponding to Çukurova Region (A) and Gediz Basin (B), respectively.

**Figure 5.**
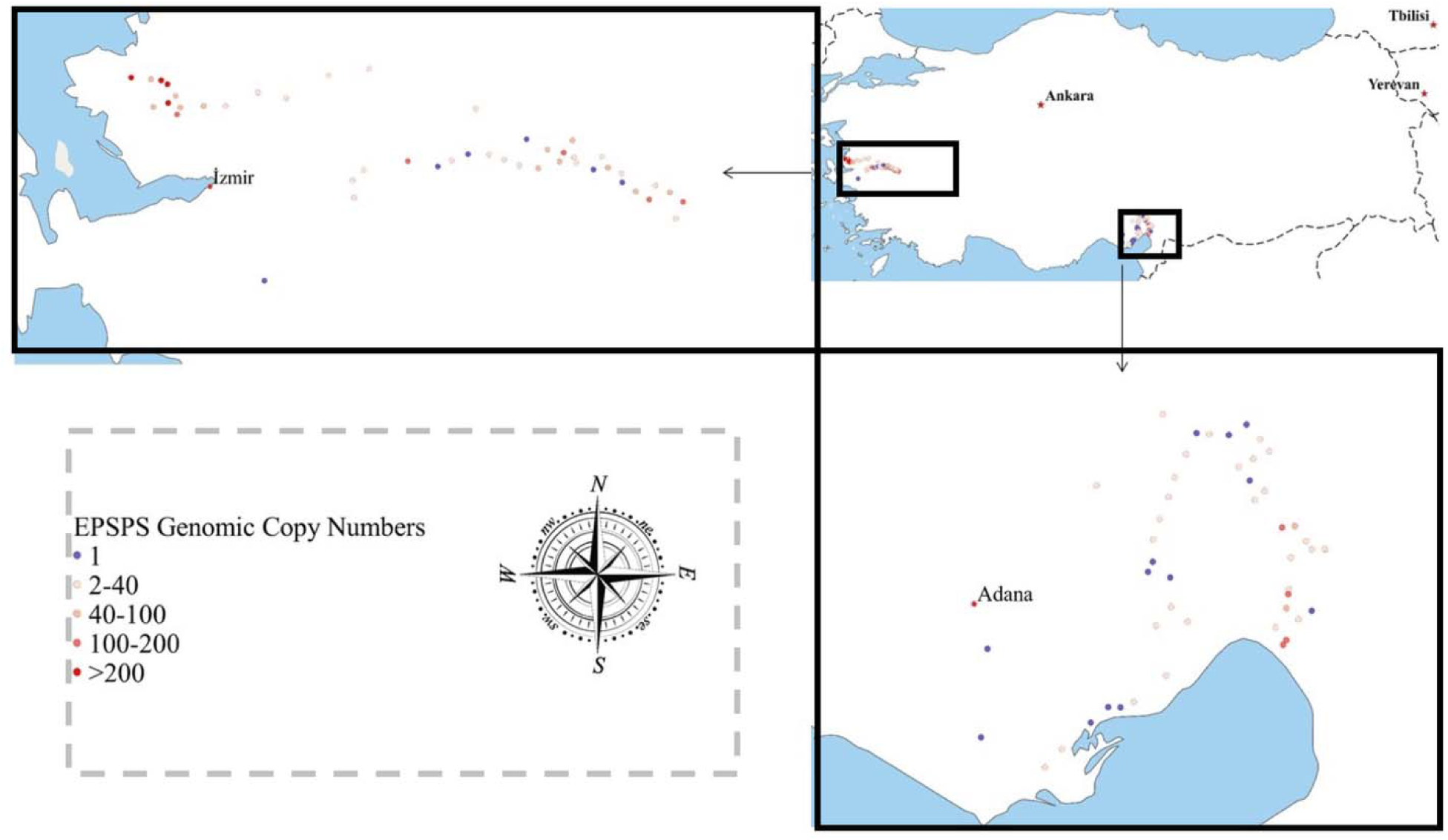
Geographic distribution relative *EPSPS* copy numbers (2^ΔCt^) in Palmer amaranth accessions across Türkiye. The main map displays the surveyed collection sites, with zoomed-in insets highlighting the Gediz Basin (western Türkiye, top left) near İzmir and the Çukurova Region (southern Türkiye, bottom right) near Adana. Colored symbols represent the specific relative *EPSPS* copy number range identified at each sampling location, as defined in the legend.

### Resistance phenotyping

Main and interaction effects of dry weight response of *A. palmeri* after application of double dose nicosulfuron and glyphosate was given in Table 3. Since no significant main and interaction effects of experimental run were observed, the data were pooled and subsequently analyzed to assess the relationship between relative dry weight and the presence of TSR mechanism.

**Table 3.** Effects of region and experimental run on the relative dry weight of *Amaranthus palmeri* after 2× herbicide application. N: Number, p: Probability value.

| Relative Dry Weight of *A. palmeri* (%) |  |  |  |
| --- | --- | --- | --- |
| Factors | N | (After 2x Nicosulfuron Application) | (After 2x Glyphosate Application) |
| Region |  | p=0.000 | p=0.000 |
| Çukurova Region (CU) | 768 | 14.28±0.759 | 8.86±1.041 |
| Gediz Basin (GE) | 768 | 5.01±0.632 | 27.93±1.350 |
| Experimental Run |  | p=0.966 | P=0.203 |
| 1 | 768 | 9.62±0.789 | 17.31±1.217 |
| 2 | 768 | 9.67±0.764 | 19.48±1.287 |
| Region*Experimental Run |  | p=0.843 | P=0.298 |

Following the application of a 2x rate of nicosulfuron to *A. palmeri* accessions collected from the Çukurova Region, surviving accessions accumulated dry biomass equivalent to no more than 44.8% that observed in the corresponding control pots. The mean relative dry weight across all accessions was 14.3% (Supplementary Table 4). In the Gediz Basin, the maximum and mean relative dry weights were 53.7% and 5.0%, respectively. Moreover, 22.9% of accessions from the Çukurova Region had relative dry weights greater than 25%, compared with 8.3% of accessions from the Gediz Basin.

Following glyphosate application at twice the recommended rate, *A. palmeri* accessions from the Çukurova Region exhibited relative dry weights of up to 107.0%, with a mean value of 8.9% relative to the untreated control (Supplementary Table 4). In the Gediz Basin, the corresponding values maximum and mean values were 97.4% and 27.9%, respectively.

Moreover, 10.4% of accessions from the Çukurova Region exhibited relative dry weights exceeding 25%, compared to 43.8% of accessions from the Gediz Basin. These results suggest that *A. palmeri* accessions from the Çukurova Region were generally more inclined to have resistance to nicosulfuron, whereas those from the Gediz Basin to glyphosate.

For analyzing dry weight response of *A. palmeri* to nicosulfuron, *ALS* mutations frequency data were used in different multivariable regression (enter) models, all investigated target-site resistance mutations (A122S, A122T, A122V, P197A, P197R, P197T, P197S, P197I, A205V, D376E, W574l, S653N) with or without region effect were simultaneously entered into the initial model (Table 4). Mutations that did not remain statistically significant after adjustment for the presence of the other mutations are not shown in the multivariable model 1, since they did not provide additional independent explanatory power. Only predictors that retained an independent association with relative dry weight were P197S and W574L. Univariable regression model results for P197S, W574L and *EPSPS* are also presented in the table. Region was incorporated to account for potential geographic confounding in one of the multivariable model 2. In the univariable regression analyses, both W574L (β = 0.398, P < 0.001) and P197S (β = 0.392, P < 0.001) mutations were significantly associated with nicosulfuron treatments, explaining 15.8% and 15.4% of the variance, respectively. In the multivariable model 1, both mutations remained independent predictors of possible resistance (W574L: β = 0.366, P < 0.001; P197S: β = 0.360, P < 0.001), increasing the explained variance to 32.9%. Inclusion of region did not significantly improve the multivariable model 2 (P = 0.549). Regional differences in herbicide response were evident descriptively, but after accounting for TSR markers, region itself did not provide significant additional explanatory power. In the interaction model 3, P197S was not significantly associated with relative dry weight after accounting for W574L and the P197S × W574L interaction term (β = 0.300, B = 0.170, 95% CI: −0.017 to 0.358, P = 0.075).

**Table 4.** Univariable and multivariable linear regression analyses of genetic herbicide-resistance markers associated with relative dry weight in *Amaranthus palmeri*.

| Dependent Variable | Model | Predictor | Standardized $\beta$ | B (95% CI) | P value | Model R<sup>2</sup> | Adj. R<sup>2</sup> |
| --- | --- | --- | --- | --- | --- | --- | --- |
| **Relative Dry Weight of *A. palmeri*** | Univariable | W574L | 0.398 | 0.292 (0.154–0.430) | <0.001 | 0.158 | 0.149 |
|  | Univariable | P197S | 0.392 | 0.223 (0.116–0.330) | <0.001 | 0.154 | 0.145 |
|  | Multivariable Model 1* | W574L | 0.366 | 0.269 (0.132–0.405) | <0.001 | 0.329 | 0.241 |
|  |  | P197S | 0.360 | 0.205 (0.096–0.313) | <0.001 |  |  |
|  | Multivariable Model 2* | W574L | 0.389 | 0.285 (0.138–0.433) | <0.001 | 0.332 | 0.235 |
|  |  | P197S | 0.419 | 0.239 (0.082–0.395) | 0.003 |  |  |
|  |  | Region | 0.104 | 2.880 (–6.651–12.411) | 0.549 |  |  |
|  | Interaction Model 3* | P197S | 0.300 | 0.170 (–0.017–0.358) | 0.075 | 0.346 | 0.233 |
|  |  | W574L | 0.319 | 0.234 (0.067–0.401) | 0.007 |  |  |
|  |  | S653N | 0.130 | 0.240 (–0.017–0.277) | 0.082 |  |  |
|  |  | P197S*W574L | 0.222 | 0.006 (–0.003–0.014) | 0.189 |  |  |
|  |  | P197S*W574L*S653N | –0.020 | –2.99x10<sup>–5</sup> (0.000–0.000) | 0.856 |  |  |
|  | Univariable | EPSPS | 0.672 | 0.274 (0.212–0.336) | <0.001 | 0.452 | 0.446 |
|  | Multivariable Model 4 | EPSPS | 0.637 | 0.260 (0.195–0.325) | <0.001 | 0.463 | 0.451 |
|  |  | Region | 0.109 | 6.532 (–3.073–16.136) | 0.180 |  |  |

\*Predictors were retained in the multivariable models based on the prespecified model structure and/or statistical significance in the corresponding univariable analyses. Multivariable model 1 includes the effects of all ALS mutation frequencies. Multivariable model 2 includes the effects of all ALS mutation frequencies with region. Interaction model 3 includes the effects of all ALS mutation frequencies with region, P197S\*W574L and P197S\*W574L\*S653N interactions. Multivariable model 4 includes relative *EPSPS* genomic copy number and region.

W574L remained significantly and positively associated with relative dry weight (β = 0.319, B = 0.234, 95% CI: 0.067–0.401, P = 0.007). S653N was not significantly associated with relative dry weight (β = 0.130, B = 0.240, 95% CI: −0.017 to 0.277, P = 0.082). The P197S × W574L interaction term was not statistically significant (β = 0.222, B = 0.006, 95% CI: −0.003 to 0.014, P = 0.189), indicating no evidence of a statistically significant interaction between P197S and W574L on relative dry weight. The three-way interaction between P197S, W574L, and S653N was also not significant (β = −0.020, B = −2.99 × 10, P = 0.856). These results suggest that P197S and W574L were the mutations associated with relative dry response to nicosulfuron at the maternal-family level.

For dry weight response of *A. palmeri* to glyphosate, relative *EPSPS* copy number showed a strong positive association in both univariable (β = 0.672, R² = 0.452, P < 0.001) and multivariable analyses (β = 0.637, R² = 0.463, P < 0.001). Region was not a significant predictor (P = 0.180), indicating that relative *EPSPS* copy number was the primary determinant of dry weight response of *A. palmeri* in multivariable model 4 (Table 4).

The relatively weak correlation observed between *ALS* mutations (W574L=0.398, P197S= 0.392) and dry weight response to nicosulfuron is likely attributable to the low frequency of these mutations within some accessions and usage of different offspring of same maternal plants, resulting in variable responses among replicates in the pot experiments. Specifically, after application of nicosulfuron at twice the recommended rate, some replicates of the same accession survived whereas others did not. In contrast, glyphosate responses were highly consistent among replicates, with all plants within a given accession either survived or died following treatment. Scatter plot diagrams of relationships between resistance-associated markers and dry weight response of *A. palmeri* accessions were given in Figure 6.

**Figure 6.**
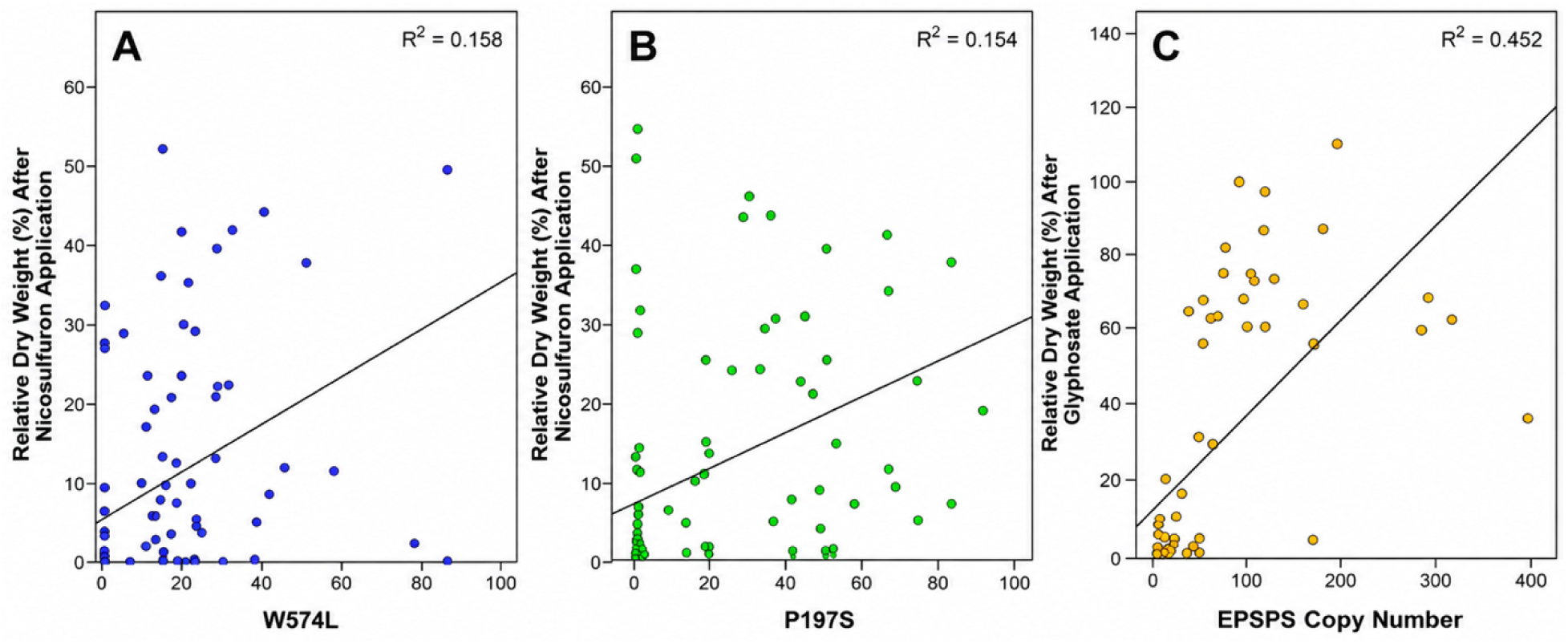
Linear relationships between relative dry weight after herbicide application and W574L mutation frequency (A), P197S mutation frequency (B), and relative *EPSPS* copy number (C) in *Amaranthus palmeri* accessions. R²: Model coefficients of determination.

## Discussion

Previous studies on glyphosate resistance in *A. palmeri* in southern Türkiye have shown an increase in relative *EPSPS* copy numbers ranging from 14.9 to 37.0 with the resistance index ranging from 2.4-7.0 (Mennan et al. 2021, Kaya Altop et al. 2025). In this study, it was determined that the relative *EPSPS* copy number did not exceed 40 in most of the accessions obtained from the Çukurova Region in southern Türkiye (87.5%), while in the Gediz Basin in western Türkiye, relative *EPSPS* genomic copy numbers were found to reach up to 397.1, and accessions with copy numbers above 40 constituted 45.83% of the total number of accessions.

The study by Kaya Altop et al. (2025) also demonstrated nicosulfuron (9.21–10.35-fold) and foramsulfuron+iodosulfuron methyl-sodium resistance (6.41–7.44-fold) in *A. palmeri* accessions in Türkiye. In the same study, the amino acid changes associated with these resistances were identified as A122V and P197R. In this study, however, different amino acid substitutions (P197S, P197I, W574L and S653N) were observed and were dominant (except for P197I) compared with those reported by Kaya Altop et al. (2025).

*ALS* mutations were more prevalent in the Çukurova Region where annual crops are cultivated (Durmuş and Yiğit 2015) and several *ALS* inhibitors are used. The Gediz Basin is an area with a high concentration of vineyards, and accessions with high relative *EPSPS* copy numbers have been found in areas where glyphosate use is intensive, such as along roadsides, irrigation canals, and in vineyards. In the Çukurova Region, the accessions with the highest relative *EPSPS* genomic copy number were found along the roadside and in olive and citrus orchards. The regional differences observed in *ALS* mutation frequencies and relative *EPSPS* genomic copy numbers are consistent with, but do not by themselves demonstrate, an effect of regional herbicide-use and cropping practices. These patterns may also reflect differences in introduction history, founder effects, genetic drift, and regional gene flow. It is unknown whether the increases in relative *EPSPS* copy numbers observed in the Gediz Basin and the Çukurova Region originate from the regions where *A. palmeri* was introduced to Türkiye. However, Gaines et al. (2010) stated that *EPSPS* gene amplification can occur in less than 7 years with repeated glyphosate use. In particular, the production pattern involving vineyards in the Gediz Basin has persisted for many years. *EPSPS* amplification can evolve rapidly under recurrent glyphosate selection, but our data cannot determine whether the observed amplification arose after introduction into Türkiye or was already present in founding material. Although dose-response studies were not conducted in this study, it has been repeatedly demonstrated by others (Culpepper et al. 2006; Singh et al. 2018; Martins et al. 2026) that accessions showing high relative *EPSPS* copy numbers will also be classified as glyphosate-resistant. Therefore, it is necessary to test herbicides that could be alternatives to glyphosate in *A. palmeri* management and to determine whether resistance to these herbicides is also developed.

The current situation indicates that *ALS* inhibitors will likely fail to control *A. palmeri*, especially in the Çukurova Region. Çatıkkaş (2024) reported that in his pot and field studies using mixed Turkish populations of *A. palmeri,* no herbicide provided sufficient control in sunflower, tomato, and cotton. In corn, however, dimethenamid-p + terbuthylazine, S-metolachlor + terbuthylazine, and isoxaflutole + thiencarbazone-methyl + cyprosulfamide were found to be effective against *A. palmeri* before emergence. Post-emergence usage of dimethenamide-p + terbuthylazine, and terbuthylazine + mesotrione also showed over 90% control efficacy. Since *A. palmeri* has been reported to acquire resistance to many of these herbicides (Souza et al. 2025; Heap 2026), it is necessary to determine whether it acquires resistance to other herbicide mechanisms of action at the population level.

Singh et al. (2019) reported that resistance to *ALS* inhibitors in *A. palmeri* in the USA was primarily due to amino acid substitutions at positions A122, P197, W574, and S653; with the A122 or S653 mutations primarily conferring resistance to the imidazolinone (IMI) *ALS* inhibitors with low-level cross-resistance to sulfonylureas (SU) (Bernasconi et al. 1995; Devine and Eberlein 1997). The P197 substitution has been shown to confer high levels of resistance to SU herbicides (Guttieri et al. 1992; Nakka et al. 2017) with low risk of IMI cross-resistance, whereas the W574 substitution confers resistance to a broad range of IMI, SU, and triazolopyrimidine (TP) herbicides. Molin et al. (2016) specifically reported imazethapyr, nicosulfuron, pyrithiobac, and trifloxysulfuron resistance resulting from the W574L mutation in spiny amaranth (*Amaranthus spinosus* L.) x *A. palmeri* hybrids. Larran et al. (2017) also associated A122, W574, and S653 mutations in *A. palmeri* with resistance to chlorimuron-ethyl, diclosulam, and imazethapyr. Palmieri et al. (2022b) found that the D376E, A205V, and A122S substitutions in *A. palmeri* conferred cross-resistance to the most used chemical families in HRAC Group 2 and confirmed that the A205V substitution conferred resistance to herbicides in the triazolopyrimidine family. In this study, the dry weight responses of *A. palmeri* accessions to nicosulfuron were positively correlated with the P197S and W574L substitutions. Although the S653 substitution was observed at a high frequency, its lack of association with nicosulfuron response suggests that the studied *A. palmeri* accessions may exhibit resistance to other herbicides, particularly those in the IMI chemical family.

The contrasting target site mechanism profiles between the Çukurova Region and the Gediz Basin likely reflect a combination of factors rather than local herbicide use alone. As an invasive, dioecious species with high dispersal and reproductive capacity, *A. palmeri* can rapidly establish populations shaped by both historical and contemporary evolutionary processes (Borgato et al. 2025). The observed regional differences in *ALS* mutations and relative *EPSPS* copy number may partly originate from independent introduction events involving genetically distinct source populations, potentially carrying pre-existing resistance traits. In addition, founder effects and genetic drift during establishment could have amplified initial differences between regions. Gene flow through pollen and human-mediated seed movement may further contribute to the spread and mixing of resistance alleles across regions, although its relative contribution cannot be quantified in this study. Local herbicide selection likely also plays a role, as *ALS* inhibitor use in Çukurova and glyphosate-intensive systems in Gediz Basin are consistent with the observed target site resistance mechanisms. However, these associations are correlative, the present data do not allow a direct casual inference.

The high frequency of the S653N mutation in the Çukurova Region despite its lack of association with nicosulfuron response suggests that its distribution may reflect historical or concurrent selection by imidazoline herbicides rather than nicosulfuron selection alone. Variation among progeny within maternal families and the potential contribution of unmeasured target-site or non-target-site resistance mechanisms may also contribute to the observed phenotype-marker discrepancies. These findings highlight the need for further investigation of the broader resistance history and genetic background of *A. palmeri* populations in Türkiye.

## Conclusion

It has been concluded that the P197S, W574L, and S653N mutations are more prevalent in *A. palmeri* accessions from Türkiye’s Çukurova region—suggesting a higher likelihood of encountering cases of resistance to *ALS* inhibitors—whereas relative EPSPS copy number levels are higher in the Gediz Basin, indicating a greater probability of encountering glyphosate resistance. Therefore, it is necessary to confirm herbicide resistance in these regions through dose-response experiments and to develop alternative control strategies. Overall, the results suggest that regional target site resistance mechanisms patterns in *A. palmeri* in Türkiye most likely arise from the interaction of introduction history, demographic processes, gene flow, and local selection rather than any single mechanism.

## Acknowledgments

Thanks to the undergraduate and graduate students of Patterson Lab/MSU who participated in the pot and laboratory studies and the colleagues and friends who helped to collect and clean Palmer amaranth seeds in Türkiye.

## Funding

This research received no specific grant from any funding agency, commercial or not-for-profit sectors. However, we would like to thank Fulbright Türkiye for covering the living expenses of the Turkish researcher as a visiting scholar, thus enabling her to work in the laboratories of Michigan State University.

## Competing Interests

The authors declare none.

## Notes

### Competing Interest Statement

The authors have declared no competing interest.

