## Supplemental Data for "Distribution of target-site resistance mechanisms to nicosulfuron and glyphosate in *Amaranthus palmeri* accessions from two different regions of Türkiye"

^1^Associate Professor, Plant Protection Department, Aydın Adnan Menderes University, Aydın, Türkiye; ^2^Postdoctoral Researcher, Department of Plant, Soil and Microbial Sciences, Michigan State University, East Lansing, MI, USA; ^3^Graduate Student, Department of Plant, Soil and Microbial Sciences, Michigan State University, East Lansing, MI, USA; ^4^Associate Professor, Department of Plant, Soil and Microbial Sciences, Michigan State University, East Lansing, MI, USA

***Authors for correspondence:**

Eric L. Patterson,

**Supplementary Material**

Supplementary Table 1. Seed collection sites of *Amaranthus palmeri* biotypes used in this study, including sample codes, geographic coordinates, and associated field or habitat type.

| Sample^a^ | Coordinates | Location | Sample^a^ | Coordinates | Location |
| --- | --- | --- | --- | --- | --- |
| CU1 | N 37°14'18.348"  E 35°38'13.6428" | Peanut | GE1 | N 38°27'4.52780"  E 27°34'12.80280" | Corn |
| CU2 | N 37°23'00.7836"  E 35°48'37.5732" | Roadside | GE2 | N 38°24'44.42940"  E 27°34'28.00650" | Peach |
| CU3 | N 37°20'43.2168"  E 35°53'55.2876" | Pomegranate | GE3 | N 38°28'30.27430"  E 27°36'8.76900" | Roadside |
| CU4 | N 37°20'35.1672"  E 35°55'52.752" | Corn | GE4 | N 38°29'44.04080"  E 27°43'59.49670" | Roadside |
| CU5 | N 37°20'29.0004"  E 35°58'59.502" | Peanut | GE5 | N 38°29'0.88010"  E 27°49'17.81340" | Roadside |
| CU6 | N 37°21'45.36"  E 36°01'44.094" | Roadside | GE6 | N 38°29'49.10100"  E 27°51'43.32720" | Tomato |
| CU7 | N 37°18'08.2152"  E 35°52'15.5172" | Corn | GE7 | N 38°30'39.84290"  E 27°54'37.51930" | Roadside |
| CU8 | N 37°15'16.5708"  E 35°50'27.2724" | Corn | GE8 | N 38°30'37.56240"  E 27°58'22.97880" | Vineyard |
| CU9 | N 37°12'54.7452  E 35°49'29.7156" | Peanut | GE9 | N 38°29'56.70510"  E 28°0'59.62120" | Vineyard |
| CU10 | N 37°10'13.3716"  E 35°48'09.342" | Corn | GE10 | N 38°29'9.09170"  E 28°3'40.69660" | Roadside |
| CU11 | N 37°07'41.3256"  E 35°47'09.2184" | Peanut | GE11 | N 38°28'45.47210"  E 28°7'4.19760" | Roadside |
| CU12 | N 37°04'59.2464"  E 35°47'03.5412" | Roadside | GE12 | N 38°29'41.34820"  E 28°10'51.09140" | Roadside |
| CU13 | N 37°03'43.8768"  E 35°46'18.7716" | Peach | GE13 | N 38°29'30.97840"  E 28°13'50.20650" | Roadside |
| CU14 | N 37°03'02.556"  E 35°49'46.704" | Cotton | GE14 | N 38°28'35.90690"  E 28°16'50.04340" | Vineyard |
| CU15 | N 36°59'27.4884"  E 35°49'57.0252" | Cotton | GE16 | N 38°26'50.67880"  E 28°21'55.13090" | Roadside |
| CU16 | N 36°57'09.6948"  E 35°47'33.4464" | Cotton | GE17 | N 38°25'36.19900"  E 28°24'18.05660" | Vineyard |
| CU17 | N 36°57'37.6704"  E 35°52'33.2076" | Corn | GE18 | N 38°24'30.55480"  E 28°26'40.54280" | Vineyard |
| CU19 | N 36°51'01.404"  E 35°49'08.3784" | Roadside | GE20 | N 38°21'58.73820"  E 28°31'22.66610" | Vineyard |
| CU20 | N 36°47'47.6808"  E 35°44'04.9272" | Soybean | GE21 | N 38°24'11.53890"  E 28°32'42.32710" | Roadside |
| CU21 | N 36°47'06.0576"  E 35°42'00.2736" | Soybean | GE22 | N 38°25'29.57740"  E 28°30'20.52810" | Vineyard |
| CU22 | N 36°47'08.1384"  E 35°40'05.592" | Citrus | GE23 | N 38°26'25.08490"  E 28°27'44.75520" | Vineyard |
| CU23 | N 36°45'12.6396"  E 35°37'21.4536" | Roadside | GE24 | N 38°26'25.42520"  E 28°27'42.97760" | Vineyard |
| CU24 | N 36°41'57.2604  E 35°32'51.8388" | Roadside | GE25 | N 38°28'3.73690"  E 28°21'46.23830" | Olive |
| CU25 | N 36°39'43.6716"  E 35°30'11.232" | Citrus | GE26 | N 38°28'50.53300"  E 28°19'24.37310" | Vineyard |
| CU26 | N 36°43'24.6684"  E 35°20'10.536" | Roadside | GE27 | N 38°30'19.21870"  E 28°18'19.17960" | Corn |
| CU29 | N 36°54'17.8236"  E 35°21'10.4292" | Roadside | GE28 | N 38°30'16.73420"  E 28°13'22.54520" | Roadside |
| CU30 | N 37°06'28.8576"  E 36°14'05.7948" | Corn | GE29 | N 38°30'53.14170"  E 28°11'33.86430" | Corn |
| CU31 | N 37°06'30.2508"  E 36°11'59.7228" | Corn | GE30 | N 38°32'31.61990"  E 28°13'7.21250" | Vineyard |
| CU33 | N 37°09'21.33"  E 36°09'19.224" | Soybean | GE31 | N 38°31'17.74920"  E 28°8'41.14490" | Vineyard |
| CU34 | N 37°09'11.0628"  E 36°07'19.4196" | Roadside | GE32 | N 38°32'41.91730"  E 28°5'0.35150" | Olive |
| CU35 | N 37°12'32.7564"  E 36°03'02.8008" | Corn | GE33 | N 38°36'49.74760"  E 27°56'4.03760" | Pepper |
| CU36 | N 37°13'38.73"  E 36°04'34.0572" | Soybean | GE35 | N 38°42'14.19630"  E 27°37'5.39040" | Corn |
| CU37 | N 37°14'54.6288"  E 36°02'14.6076" | Peanut | GE36 | N 38°42'14.41030"  E 27°37'5.24680" | Vineyard |
| CU38 | N 37°16'35.1228"  E 36°00'32.5836" | Corn | GE37 | N 38°41'22.83900"  E 27°29'57.10570" | Roadside |
| CU39 | N 37°17'34.8432"  E 36°02'48.642" | Peanut | GE38-1 | N 38°38'17.90030"  E 27°22'26.15240" | Vineyard |
| CU40 | N 37°18'30.0456"  E 36°05'15.9612" | Peanut | GE38-2 | N 38°39'2.84980"  E 27°17'27.46690" | Tomato |
| CU41 | N 37°19'58.9368"  E 36°03'51.2424" | Roadside | GE39 | N 38°37'10.64850"  E 27°11'44.92420" | Roadside |
| CU42 | N 37°07'29.4132"  E 36°11'06.5292" | Peanut | GE40 | N 38°37'13.05690"  E 27°7'51.87960" | Roadside |
| CU43 | N 37°05'28.4136"  E 36°08'40.938" | Soybean | GE41 | N 38°36'3.15310"  E 27°3'8.61750" | Irrigation Channel Side |
| CU44 | N 37°06'34.6212"  E 36°14'00.4236" | Roadside | GE42 | N 38°37'0.09970"  E 27°3'44.01640" | Roadside |
| CU45 | N 37°01'36.0264"  E 36°08'21.7608" | Okra | GE43 | N 38°38'33.77160"  E 27°2'54.50150" | Vineyard |
| CU46 | N 37°01'00.3396"  E 36°08'17.2464" | Citrus | GE44 | N 38°40'8.59450"  E 27°1'25.85470" | Irrigation Channel Side |
| CU47 | N 36°59'16.6956"  E 36°08'00.2904" | Olive | GE45 | N 38°40'39.58710"  E 27°0'20.47640" | Plum |
| CU49 | N 36°56'53.7576"  E 36°06'15.93" | Citrus | GE46 | N 38°40'48.26810"  E 26°58'32.65380" | Roadside |
| CU50 | N 36°54'47.304"  E 36°07'29.2476" | Citrus | GE48-1 | N 38°41'3.15490"  E 26°55'2.52630" | Roadside |
| CU51 | N 36°55'22.44"  E 36°07'59.4228" | Citrus | GE48-2 | N 38°37'4.48540"  E 26°58'56.46560" | Vineyard |
| CU52 | N 36°57'55.9044"  E 36°09'52.4304" | Citrus | GE49 | N 38°37'36.08820"  E 27°1'33.76710" | Green Pea |
| CU53 | N 36°58'58.5732"  E 36°11'57.2604" | Citrus | GE50 | N 38°13'28.12120"  E 27°18'35.60480" | Roadside |

^a^The abbreviation CU refers to the Çukurova Region (southern part) of Türkiye, and the abbreviation GE refers to the Gediz Basin (western part) of Türkiye.

Supplementary Table 2. Mean frequency of *ALS* target-site mutations (>5%) of accessions from the Çukurova Region (CU) and Gediz Basin (GE) of Türkiye.

| Biotypes | P197S | P197I | W574L | S653N |  | Biotypes | P197S | P197I | W574L | S653N |
| --- | --- | --- | --- | --- | --- | --- | --- | --- | --- | --- |
| CU-01 | 18.00 | 0.00 | 12.00 | 51.60 |  | GE-01 | 0.50 | 0.00 | 77.40 | 0.00 |
| CU-02 | 48.20 | 0.00 | 24.40 | 24.10 |  | GE-02 | 0.30 | 0.00 | 39.90 | 0.20 |
| CU-03 | 0.60 | 0.00 | 24.70 | 73.20 |  | GE-03 | 17.50 | 0.00 | 0.10 | 10.10 |
| CU-04 | 0.00 | 0.00 | 22.90 | 76.40 |  | GE-04 | 0.60 | 0.00 | 11.60 | 0.00 |
| CU-05 | 1.20 | 0.00 | 21.80 | 77.20 |  | GE-05 | 0.50 | 0.10 | 0.30 | 0.20 |
| CU-06 | 67.00 | 0.00 | 15.50 | 17.80 |  | GE-06 | 0.30 | 0.00 | 0.00 | 0.30 |
| CU-07 | 89.30 | 0.00 | 11.60 | 0.00 |  | GE-07 | 0.50 | 0.00 | 0.10 | 0.10 |
| CU-08 | 28.20 | 0.00 | 41.80 | 0.20 |  | GE-08 | 0.90 | 0.00 | 0.00 | 16.00 |
| CU-09 | 35.10 | 0.00 | 5.30 | 59.10 |  | GE-09 | 2.20 | 0.60 | 0.00 | 23.90 |
| CU-10 | 81.00 | 0.00 | 13.30 | 1.00 |  | GE-10 | 0.50 | 0.00 | 0.10 | 0.10 |
| CU-11 | 18.90 | 0.00 | 48.10 | 17.20 |  | GE-11 | 0.00 | 0.00 | 0.00 | 0.30 |
| CU-12 | 72.30 | 0.00 | 30.00 | 0.60 |  | GE-12 | 0.60 | 0.10 | 0.00 | 0.40 |
| CU-13 | 48.10 | 0.00 | 21.10 | 28.10 |  | GE-13 | 1.10 | 0.00 | 16.40 | 0.20 |
| CU-14 | 65.40 | 0.20 | 10.30 | 9.90 |  | GE-14 | 1.30 | 0.00 | 0.00 | 0.00 |
| CU-15 | 24.50 | 0.00 | 30.60 | 22.10 |  | GE-16 | 0.90 | 0.00 | 0.00 | 0.00 |
| CU-16 | 44.70 | 0.20 | 13.80 | 37.40 |  | GE-17 | 0.50 | 0.20 | 0.10 | 33.20 |
| CU-17 | 47.30 | 0.00 | 52.00 | 0.10 |  | GE-18 | 0.70 | 0.00 | 0.10 | 0.30 |
| CU-19 | 0.80 | 0.00 | 25.20 | 72.90 |  | GE-20 | 0.60 | 0.00 | 0.00 | 0.30 |
| CU-20 | 33.50 | 0.00 | 33.90 | 33.10 |  | GE-21 | 0.40 | 0.00 | 0.10 | 0.10 |
| CU-21 | 1.00 | 0.00 | 19.80 | 13.80 |  | GE-22 | 0.60 | 0.00 | 0.10 | 0.20 |
| CU-22 | 15.20 | 0.00 | 43.60 | 32.20 |  | GE-23 | 1.30 | 0.00 | 0.00 | 0.10 |
| CU-23 | 46.90 | 0.00 | 14.30 | 33.10 |  | GE-24 | 0.70 | 0.00 | 0.00 | 0.00 |
| CU-24 | 39.20 | 0.00 | 0.00 | 59.80 |  | GE-25 | 0.60 | 0.00 | 20.30 | 14.20 |
| CU-25 | 65.10 | 0.00 | 0.00 | 30.50 |  | GE-26 | 0.50 | 0.00 | 14.70 | 0.00 |
| CU-26 | 0.80 | 0.00 | 25.20 | 76.30 |  | GE-27 | 0.20 | 0.10 | 0.10 | 0.10 |
| CU-29 | 42.60 | 0.00 | 24.70 | 25.50 |  | GE-28 | 0.40 | 0.10 | 23.60 | 5.40 |
| CU-30 | 0.50 | 0.00 | 0.00 | 98.90 |  | GE-29 | 0.00 | 0.00 | 60.30 | 0.10 |
| CU-31 | 46.60 | 0.00 | 19.90 | 32.80 |  | GE-30 | 0.20 | 0.00 | 0.00 | 15.40 |
| CU-33 | 39.60 | 0.10 | 16.20 | 16.90 |  | GE-31 | 0.10 | 0.00 | 0.00 | 0.40 |
| CU-34 | 50.10 | 0.00 | 0.10 | 5.70 |  | GE-32 | 0.70 | 0.00 | 0.00 | 0.00 |
| CU-35 | 50.90 | 0.10 | 29.90 | 0.10 |  | GE-33 | 1.00 | 0.10 | 0.00 | 0.20 |
| CU-36 | 8.40 | 0.20 | 40.50 | 0.20 |  | GE-35 | 80.80 | 0.00 | 15.30 | 0.30 |
| CU-37 | 72.80 | 0.10 | 0.10 | 34.40 |  | GE-36 | 0.90 | 34.60 | 22.20 | 20.70 |
| CU-38 | 34.50 | 0.00 | 26.60 | 40.30 |  | GE-37 | 0.80 | 0.00 | 0.00 | 0.20 |
| CU-39 | 32.40 | 0.30 | 0.00 | 66.00 |  | GE-38/1 | 0.30 | 0.00 | 7.20 | 0.10 |
| CU-40 | 13.00 | 0.00 | 86.70 | 0.20 |  | GE-38/2 | 1.20 | 0.10 | 16.90 | 0.00 |
| CU-41 | 18.20 | 0.20 | 16.00 | 64.70 |  | GE-39 | 0.50 | 0.00 | 0.20 | 36.90 |
| CU-42 | 0.20 | 0.10 | 0.00 | 65.60 |  | GE-40 | 41.60 | 0.00 | 18.40 | 0.10 |
| CU-43 | 26.80 | 0.20 | 21.10 | 19.30 |  | GE-41 | 1.10 | 0.00 | 0.00 | 0.10 |
| CU-44 | 18.80 | 0.10 | 0.20 | 0.10 |  | GE-42 | 0.60 | 0.00 | 0.00 | 0.30 |
| CU-45 | 12.70 | 0.10 | 18.40 | 59.10 |  | GE-43 | 1.30 | 0.00 | 0.00 | 0.20 |
| CU-46 | 1.70 | 0.00 | 0.00 | 99.00 |  | GE-44 | 0.70 | 0.00 | 0.00 | 54.90 |
| CU-47 | 0.60 | 0.20 | 32.10 | 37.00 |  | GE-45 | 0.40 | 0.00 | 0.20 | 37.70 |
| CU-49 | 0.50 | 0.00 | 0.20 | 41.70 |  | GE-46 | 0.50 | 0.10 | 0.10 | 40.70 |
| CU-50 | 31.30 | 0.00 | 33.20 | 30.60 |  | GE-48-1 | 0.90 | 0.00 | 0.00 | 39.90 |
| CU-51 | 17.90 | 0.10 | 0.00 | 16.00 |  | GE-48-2 | 0.20 | 0.00 | 0.20 | 0.30 |
| CU-52 | 55.80 | 0.00 | 14.10 | 20.60 |  | GE-49 | 0.50 | 0.10 | 0.20 | 0.10 |
| CU-53 | 64.60 | 0.00 | 30.30 | 0.00 |  | GE-50 | 0.10 | 0.00 | 84.10 | 18.00 |

Abbr. CU=Çukurova Region, GE= Gediz Basin, P=Pro (Proline), S=Ser (Serine), I=Ile (Isoleucine), W=Trp (Tryptophan), L=Leu (Leucine), N= Asn (Asparagine)

Supplementary Table 3. Relative EPSPS genomic copy number variation (CNV) in *Amaranthus palmeri* biotypes, expressed relative to the single-copy reference gene A36.

| Biotype | CNV | SE | Biotype | CNV | SE |
| --- | --- | --- | --- | --- | --- |
| CU-01 | 1.69 | 0.14 | GE-01 | 1.66 | 0.10 |
| CU-02 | 1.63 | 0.07 | GE-02 | 15.46 | 14.13 |
| CU-03 | 1.04 | 0.24 | GE-03 | 1.60 | 0.18 |
| CU-04 | 1.68 | 0.13 | GE-04 | 111.59 | 45.30 |
| CU-05 | 1.47 | 0.53 | GE-05 | 1.41 | 0.23 |
| CU-06 | 1.10 | 0.30 | GE-06 | 1.73 | 0.21 |
| CU-07 | 1.63 | 0.15 | GE-07 | 1.48 | 0.13 |
| CU-08 | 1.72 | 0.10 | GE-08 | 31.36 | 23.94 |
| CU-09 | 10.94 | 9.46 | GE-09 | 25.79 | 12.79 |
| CU-10 | 1.55 | 0.08 | GE-10 | 11.83 | 10.65 |
| CU-11 | 1.90 | 0.36 | GE-11 | 43.75 | 21.30 |
| CU-12 | 1.44 | 0.03 | GE-12 | 47.77 | 12.83 |
| CU-13 | 1.18 | 0.29 | GE-13 | 17.94 | 15.63 |
| CU-14 | 1.34 | 0.36 | GE-14 | 1.06 | 0.22 |
| CU-15 | 1.61 | 0.29 | GE-16 | 1.43 | 0.12 |
| CU-16 | 1.68 | 0.10 | GE-17 | 82.88 | 19.10 |
| CU-17 | 1.52 | 0.26 | GE-18 | 175.57 | 66.74 |
| CU-19 | 1.61 | 0.18 | GE-20 | 17.44 | 15.20 |
| CU-20 | 2.03 | 0.47 | GE-21 | 123.76 | 41.36 |
| CU-21 | 1.35 | 0.31 | GE-22 | 89.80 | 37.00 |
| CU-22 | 1.33 | 0.35 | GE-23 | 100.86 | 20.32 |
| CU-23 | 1.42 | 0.03 | GE-24 | 37.50 | 36.47 |
| CU-24 | 1.62 | 0.11 | GE-25 | 3.10 | 2.01 |
| CU-25 | 1.65 | 0.09 | GE-26 | 43.58 | 22.97 |
| CU-26 | 1.20 | 0.14 | GE-27 | 20.26 | 18.71 |
| CU-29 | 1.31 | 0.37 | GE-28 | 8.16 | 4.78 |
| CU-30 | 1.71 | 0.14 | GE-29 | 162.56 | 81.11 |
| CU-31 | 1.65 | 0.23 | GE-30 | 69.77 | 16.96 |
| CU-33 | 72.09 | 36.53 | GE-31 | 62.63 | 27.12 |
| CU-34 | 184.20 | 44.70 | GE-32 | 0.70 | 0.16 |
| CU-35 | 1.76 | 0.25 | GE-33 | 1.52 | 0.24 |
| CU-36 | 1.66 | 0.05 | GE-35 | 6.85 | 5.51 |
| CU-37 | 1.48 | 0.31 | GE-36 | 32.22 | 14.87 |
| CU-38 | 1.87 | 0.21 | GE-37 | 8.55 | 7.04 |
| CU-39 | 1.83 | 0.08 | GE-38/1 | 1.74 | 0.25 |
| CU-40 | 1.85 | 0.01 | GE-38/2 | 8.53 | 6.59 |
| CU-41 | 1.76 | 0.09 | GE-39 | 2.21 | 0.19 |
| CU-42 | 9.22 | 8.21 | GE-40 | 56.08 | 9.54 |
| CU-43 | 30.86 | 29.20 | GE-41 | 152.02 | 51.32 |
| CU-44 | 1.54 | 0.22 | GE-42 | 98.19 | 59.74 |
| CU-45 | 9.19 | 7.28 | GE-43 | 42.37 | 25.12 |
| CU-46 | 160.34 | 134.36 | GE-44 | 397.10 | 377.60 |
| CU-47 | 57.91 | 52.66 | GE-45 | 281.20 | 178.37 |
| CU-49 | 1.96 | 0.06 | GE-46 | 48.61 | 14.10 |
| CU-50 | 111.79 | 19.31 | GE-48-1 | 274.70 | 202.61 |
| CU-51 | 109.92 | 15.58 | GE-48-2 | 93.69 | 44.72 |
| CU-52 | 1.99 | 0.16 | GE-49 | 312.54 | 257.11 |
| CU-53 | 1.03 | 0.24 | GE-50 | 0.95 | 0.26 |

Abbr. CU=Çukurova Region, GE= Gediz Basin, SE= standard error of the mean, CNV= Number of copies of EPSPS relative to *A36*

Supplementary Table 4. Relative dry weight (%) of *Amaranthus palmeri* biotypes compared with untreated controls following application of nicosulfuron or glyphosate at 2× the recommended rate.

|  | Dry weight relative to control (%) | | | |  | Dry weight relative to control (%) | | | |
| --- | --- | --- | --- | --- | --- | --- | --- | --- | --- |
| Biotype | Nicosulfuron | SE | Glyphosate | SE | Biotype | Nicosulfuron | SE | Glyphosate | SE |
| CU-01 | 23.93 | 6.26 | 0.02 | 0.00 | GE-01 | 2.41 | 1.08 | 0.01 | 0.00 |
| CU-02 | 0.37 | 0.24 | 0.01 | 0.00 | GE-02 | 0.45 | 0.35 | 0.40 | 0.39 |
| CU-03 | 0.14 | 0.13 | 0.01 | 0.00 | GE-03 | 9.54 | 2.22 | 0.01 | 0.00 |
| CU-04 | 35.74 | 10.56 | 0.01 | 0.00 | GE-04 | 2.04 | 0.64 | 56.32 | 7.51 |
| CU-05 | 30.40 | 8.99 | 0.01 | 0.00 | GE-05 | 0.01 | 0.00 | 0.01 | 0.00 |
| CU-06 | 8.00 | 2.86 | 0.01 | 0.00 | GE-06 | 0.01 | 0.00 | 0.01 | 0.00 |
| CU-07 | 17.42 | 6.10 | 0.01 | 0.00 | GE-07 | 0.01 | 0.00 | 0.01 | 0.00 |
| CU-08 | 44.83 | 5.77 | 0.01 | 0.00 | GE-08 | 0.01 | 0.00 | 0.01 | 0.00 |
| CU-09 | 29.27 | 5.64 | 0.01 | 0.00 | GE-09 | 0.02 | 0.00 | 14.33 | 5.67 |
| CU-10 | 5.92 | 2.69 | 0.01 | 0.00 | GE-10 | 0.02 | 0.00 | 0.96 | 0.65 |
| CU-11 | 12.16 | 2.92 | 0.01 | 0.00 | GE-11 | 0.03 | 0.02 | 0.18 | 0.11 |
| CU-12 | 21.34 | 5.31 | 0.01 | 0.00 | GE-12 | 0.01 | 0.00 | 63.76 | 8.40 |
| CU-13 | 23.92 | 4.48 | 0.01 | 0.00 | GE-13 | 1.29 | 0.83 | 3.01 | 2.05 |
| CU-14 | 10.18 | 4.93 | 0.01 | 0.00 | GE-14 | 0.01 | 0.00 | 1.28 | 0.66 |
| CU-15 | 22.63 | 5.04 | 0.01 | 0.00 | GE-16 | 0.01 | 0.00 | 0.63 | 0.61 |
| CU-16 | 19.66 | 4.83 | 0.01 | 0.00 | GE-17 | 0.02 | 0.00 | 97.36 | 12.77 |
| CU-17 | 38.25 | 6.85 | 0.01 | 0.00 | GE-18 | 0.01 | 0.00 | 84.78 | 6.35 |
| CU-19 | 5.44 | 2.50 | 0.01 | 0.00 | GE-20 | 0.01 | 0.00 | 2.32 | 1.78 |
| CU-20 | 42.51 | 7.84 | 0.01 | 0.00 | GE-21 | 0.01 | 0.00 | 70.13 | 8.17 |
| CU-21 | 12.78 | 6.15 | 0.01 | 0.00 | GE-22 | 0.01 | 0.00 | 64.10 | 9.37 |
| CU-22 | 8.71 | 2.48 | 0.01 | 0.00 | GE-23 | 0.01 | 0.00 | 69.46 | 8.75 |
| CU-23 | 2.89 | 0.85 | 0.01 | 0.00 | GE-24 | 0.01 | 0.00 | 1.44 | 1.04 |
| CU-24 | 6.46 | 3.28 | 0.58 | 0.43 | GE-25 | 0.19 | 0.18 | 7.54 | 3.09 |
| CU-25 | 32.81 | 5.12 | 0.02 | 0.00 | GE-26 | 53.70 | 9.58 | 3.21 | 2.83 |
| CU-26 | 4.58 | 1.70 | 0.02 | 0.00 | GE-27 | 0.01 | 0.00 | 8.52 | 3.98 |
| CU-29 | 29.55 | 10.88 | 0.02 | 0.00 | GE-28 | 10.11 | 5.35 | 0.01 | 0.00 |
| CU-30 | 0.93 | 0.38 | 0.02 | 0.00 | GE-29 | 11.70 | 4.38 | 52.13 | 7.88 |
| CU-31 | 7.58 | 3.16 | 6.64 | 4.49 | GE-30 | 0.02 | 0.00 | 71.74 | 9.19 |
| CU-33 | 0.31 | 0.18 | 78.91 | 6.89 | GE-31 | 0.01 | 0.00 | 59.26 | 6.93 |
| CU-34 | 0.61 | 0.43 | 107.01 | 9.85 | GE-32 | 0.01 | 0.00 | 0.06 | 0.05 |
| CU-35 | 13.37 | 3.43 | 0.02 | 0.00 | GE-33 | 0.01 | 0.00 | 0.01 | 0.00 |
| CU-36 | 5.12 | 1.43 | 0.02 | 0.00 | GE-35 | 36.60 | 10.31 | 0.02 | 0.00 |
| CU-37 | 3.85 | 1.13 | 0.01 | 0.00 | GE-36 | 0.01 | 0.00 | 60.64 | 6.19 |
| CU-38 | 3.73 | 1.96 | 0.01 | 0.00 | GE-37 | 0.01 | 0.00 | 0.01 | 0.00 |
| CU-39 | 27.99 | 9.11 | 0.02 | 0.00 | GE-38/1 | 0.01 | 0.00 | 0.01 | 0.00 |
| CU-40 | 0.17 | 0.11 | 0.03 | 0.00 | GE-38/2 | 9.83 | 5.75 | 3.46 | 2.64 |
| CU-41 | 13.57 | 8.85 | 0.01 | 0.00 | GE-39 | 0.01 | 0.00 | 0.01 | 0.00 |
| CU-42 | 1.41 | 0.61 | 0.01 | 0.00 | GE-40 | 21.20 | 8.93 | 58.71 | 5.17 |
| CU-43 | 42.28 | 12.59 | 0.01 | 0.00 | GE-41 | 0.01 | 0.00 | 62.86 | 8.92 |
| CU-44 | 0.74 | 0.29 | 0.01 | 0.00 | GE-42 | 0.01 | 0.00 | 71.55 | 5.32 |
| CU-45 | 3.56 | 2.25 | 18.04 | 8.08 | GE-43 | 0.01 | 0.00 | 28.85 | 5.36 |
| CU-46 | 0.57 | 0.35 | 3.00 | 2.32 | GE-44 | 0.01 | 0.00 | 34.80 | 9.14 |
| CU-47 | 0.01 | 0.00 | 26.98 | 8.57 | GE-45 | 0.01 | 0.00 | 64.37 | 7.14 |
| CU-49 | 0.17 | 0.11 | 4.13 | 2.88 | GE-46 | 3.39 | 2.52 | 52.15 | 6.95 |
| CU-50 | 22.83 | 6.19 | 95.01 | 5.68 | GE-48-1 | 0.01 | 0.00 | 55.58 | 7.81 |
| CU-51 | 0.83 | 0.57 | 84.33 | 15.15 | GE-48-2 | 0.01 | 0.00 | 56.33 | 6.80 |
| CU-52 | 5.88 | 3.01 | 0.01 | 0.00 | GE-49 | 27.42 | 9.29 | 58.13 | 4.57 |
| CU-53 | 40.12 | 5.02 | 0.01 | 0.00 | GE-50 | 50.04 | 6.68 | 0.01 | 0.00 |
| Mean | 14.28 |  | 8.86 |  | Mean | 5.01 |  | 27.93 |  |

Abbr. CU=Çukurova Region, GE= Gediz Basin, SE= standard error of the mean
